# Demographic behavior of social insect populations: the specific case of Formicidae

**DOI:** 10.64898/2026.02.04.703819

**Authors:** Laura Magnani Machado, Dulce Maria de Oliveira Gomes, Filipe José Ribeiro

## Abstract

*Monomorium pharaonis* is one of the many invasive ant species which can be found associated with the endosymbiont bacteria of the genus *Wolbachia*. The association of *M. pharaonis* ants with *Wolbachia* is still being studied but is already known for giving the colony a reproductive advantage. The present work aimed to use biodemography analysis to check the effect of *Wolbachia* in the colonies of *M. pharaonis* ants, regarding its reproductive potential and rate of fertility (fertility pace). We took advantage of Birch (1948) methods to assess the effect of the bacteria in the whole colony, using data of the Dryad project. We evaluated the mean length of the generation, its capacity to multiply and the intrinsic rate of increase (*r*) and if the presence of the bacteria favors the longevity of the colony. The results obtained in the present work confirmed our initial hypothesis that the presence of the endosymbiont bacteria *Wolbachia* increased the reproductive rates of *M. pharaonis* colonies. We also found that the employment of interdisciplinary approaches highly contributes to obtaining more accurate and quantifiable results. The application of this methodological approach, highly contributed to obtain more accurate and directly driven results. For example, colonies infected with *Wolbachia* showed higher intrinsic growth rate (*r*) and thus enlightening with a new methodological approach results already presented in previous research. This "new" methodological approach revealed itself as a new tool extendible to other ants’ colonies or even other species. The use of statistical and biodemographic formulas and the adaptation of classical demography concepts for the study of the growth and reproduction of ant colonies revealed to be very useful.

## Introduction

*Monomorium pharaonis* (Hymenoptera: Formicidae) is an ant species known as “Pharaoh’s ants”, are cosmopolitan, inhabiting buildings in the temperate region of the globe and are characterized by being a highly polygynous species that multiplies by budding (1). Its reproduction is rapid, producing new offspring every few months through crossings within the colony, in addition to its queens becoming reproductive quickly (2). Ants are eusocial organisms which live in colonies, organized in the reproductive castes, such as the queens and males, and the sterile castes, such as the workers and, in some cases, soldiers. Allocation to different castes is regulated by the ratio between the number of adult workers and the number of eggs in the colony, and not by the number of queens (3).

The reproductive cycle of the Pharaoh’s ants, such as other ant species, is summarized in Fig 1. The fertile females of the colony, which are the queens, lay eggs throughout their life, after mating with a male. The larvae phase lasts from 7 to 18 days, followed by the pre-pupae and pupae period, that vary from 9 to 19 days. The pupae then differentiate into the fertile adults (queens or male individuals) or sterile adults, known as the workers of the colony (4).

**Figure 1:**
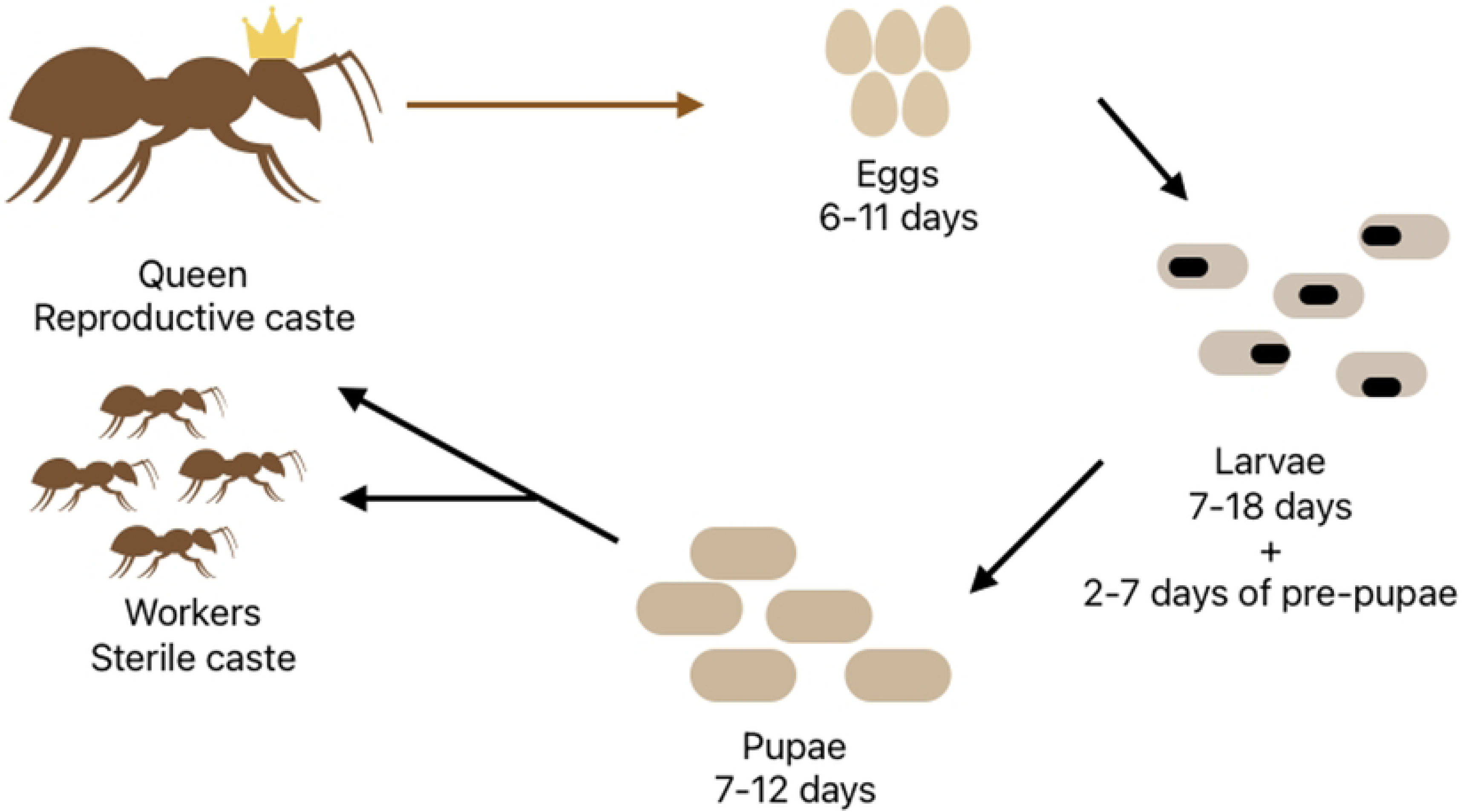
Reproductive cycle of *M. pharaonis*. Life stages with each mean time after the laying of the eggs by the queens. (Own elaboration, (4)).

The pharaoh’s ant has become an invasive species with massive pest infestations inside houses, being dependent on human activities for its survival (1) (Fig 2). Because of that, many studies regarding its colonies’ growth mechanisms have been done on the past years. Previous studies showed that pharaoh’s ants alter the allocation of resources according to its reproductive capacity and workforce of the colonies, altering the number of queens depending on the number of the eggs present at the colony (3). In some cases, workers can forage for more proteins to prioritize the growth of the colony, instead the survival in an individual level, when the colony is divided in smaller propagules (5)

**Figure 2:**
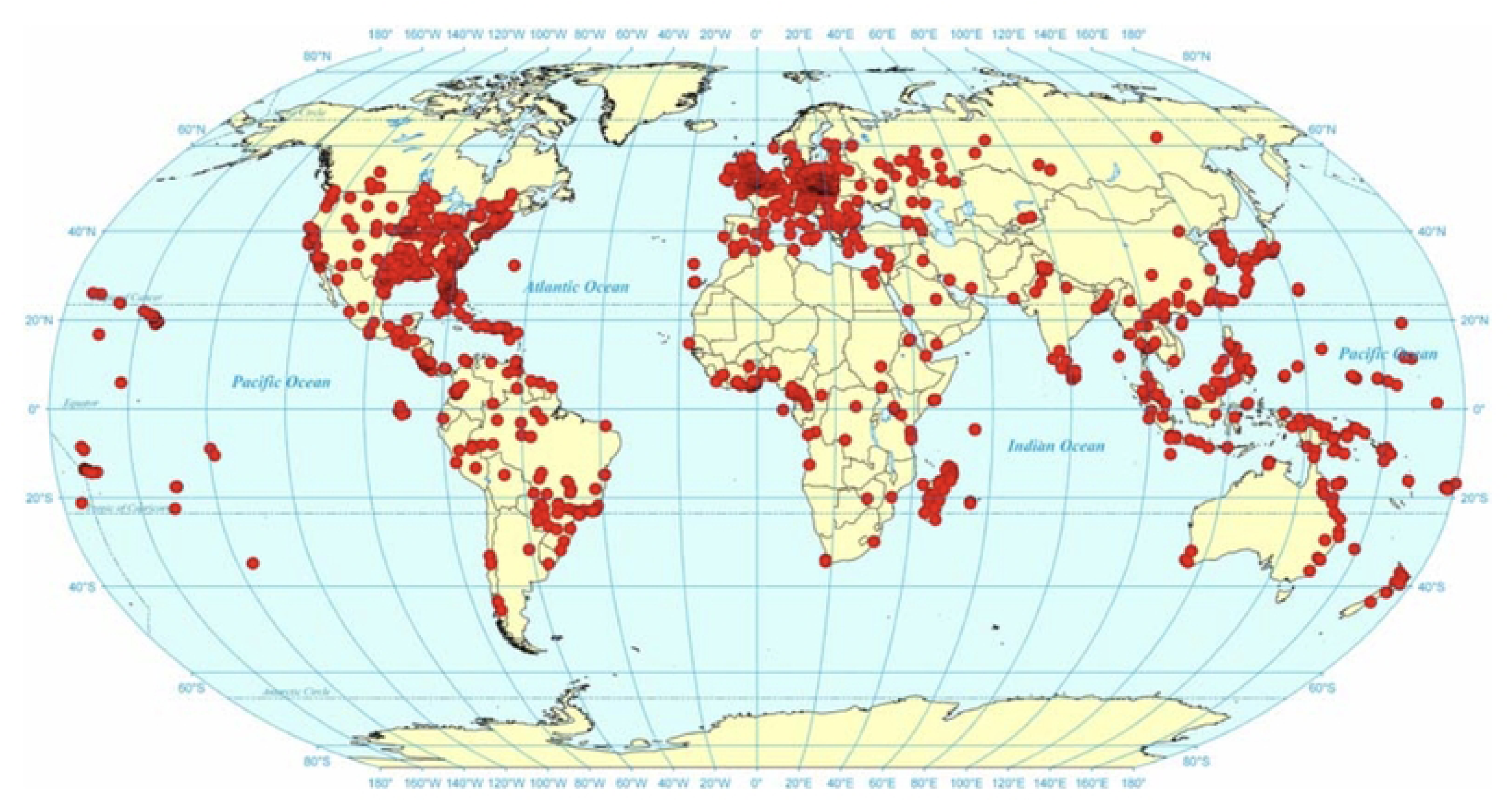
Distribution of *M. pharaonis* worldwide, according to Wetterer (6). This invasive species is a very serious urban indoor pest, mainly in hospitals, and its origin is probably Asian, but this is not well agreed within researchers.

Besides resource allocation, ants use other reproductive mechanisms related to their evolution and natural history, such as the association with endosymbiont bacteria (7–9). *Wolbachia* is a bacteria genus which, among many others, infect around 34% of the ant species known in the world and is associated with reproductive strategies (10). The association of *M. pharaonis* ants with *Wolbachia* is still being studied but is already known for giving the colony a reproductive advantage, depending on the life stage of the queen and workers, as the colony produces more reproductive individuals earlier when infected with the endosymbiont (8).

The study of insects’ populations’ growth and reproductive dynamics can benefit from the use of classic demography adapted tools. Biodemography is a subdiscipline of the classic demography, which emerged in the beginning of the century to a better understanding of the reproductive mechanisms of animal populations, going beyond the creation of specific tool for biology problems (11). Social insects are ideal for evolution studies as queens and workers have different lifespans (12). Birch (13) has developed the concept of an Intrinsic Rate of Natural Increase (*r*) for insect populations, which should be more appropriate for these studies in comparison to ‘biotic-potential’ analysis and other classical growth rates. This rate defines the intrinsic capacity of a certain population to increase, with its particulars regimes of fecundity and mortality. Besides that, this concept is not widely used in the biology studies, nor in the pharaoh’s ants-*Wolbachia* association studies.

The present work aimed to use biodemography analysis to check the effect of *Wolbachia* in the colonies of *M. pharaonis* ants, regarding its reproductive potential, with the use of biodemography concepts and rates, as well as the application of bold methodologies, such as GEE modelling and classical demography concepts to analyze the data. The database used in this work had previous studies published which stated that *Wolbachia* increased the life days of the queens and workers, as the egg laying rates (9). Having a better knowledge of the true effect *Wolbachia* has on the life span and, mainly, on the capacity it gives the colony to grow in only one generation, may help to develop more effective control methods.

### “Bug’s life”, deceleration and Plateaus in Insect colonies

Following the introductory section, this study about fecundity dynamics in insect colonies provides a unique opportunity to explore fundamental questions about fecundity, reproduction rate (pace) and survival. The key concept of this biodemographic procedure is to modulate fecundity deceleration observed at older ages by employing almost exclusively human mortality models such as the Gompertz model ((14). Nevertheless, mortality deceleration observed at advanced ages, where the exponential increase in mortality predicted by models such as the Gompertz law no longer holds (15,16) in human populations but could be useful for innovative approaches.

In humans, the force of mortality tends to a plateau, and that indicates that the rate of aging slows down or stabilizes among survivors (17). This mortality behavior is widely documented in humans and other species, but evidence also suggests that it can be also validated for social insects as ants. In this peace of research, we follow Birch’s (13) work as a first step to explore this connection.

From a theoretical perspective, Birch (13) developed a pioneering approach to study intrinsic rates increase in insect populations providing the foundation for understanding how fecundity patterns shape insect population growth and reproductive strategies. It is known that, in colonies, where the division of labor and social organization influences individuals’ survivorship, mortality plateaus may reflect both biological heterogeneity and colony-level buffering mechanisms (11,18), but it will be also possible to be observed in ant fecundity patterns?

Heterogeneity models, such as frailty distributed based ones, explain mortality deceleration because of differential survival of more robust individuals within a cohort (19). In insect colonies, more precisely for ants, this may translate into survival advantages for certain worker subgroups or castes that disproportionately contribute to late-life survival and reproductive success.

Despite that empirical analysis of insect mortality and reproduction are considered very important and informative, within this piece of research, we believe that the application of advanced demographic methods will major benefit the outcome. For example, the usually denominated hazard function estimates (fecundity rates’ hazard, in our approach) will be calculated, as well as fecundity deceleration and model comparison between Gaussian, Gamma, Weibull and Gompertz-Makeham distributions. This follows findings that hazard function estimation, survival analysis, and model comparisons between Gompertz, Gompertz–Makeham, and Weibull distributions enable precise identification of the age at which deceleration occurs (20,21).

We strongly believe thus that these approaches, when combined with life-history data from controlled colony experiments (infected vs. uninfected colonies of ants), make it possible to assess whether insect populations display longevity plateaus like those described in human demography. Understanding the onset and magnitude of fecundity deceleration in ants is not merely of theoretical interest but has practical implications for evolutionary biology and research in a broader way. Linking empirical findings to Birch’s (13) classical framework and modern biodemographic insights, we intend to contribute to broader debates on the universality of aging laws and the existence of biological limits to longevity (22,23) and fecundity in insect colonies.

## Materials and Methods

The present study investigates fecundity dynamics and late-life fecundity deceleration in ant colonies using an integrative biodemographic framework. Data was collected from controlled laboratory experiments in which queen ants were isolated and observed over their lifespans under standardized conditions. Daily records of survival and reproduction (egg counts) were maintained for each colony, allowing the construction of detailed age-specific mortality and fertility schedules.

### Database collection and description

The data was collected from the Dryad database, an open data publishing platform, (https://datadryad.org) in December of 2023. The dataset used in the present work was published by Singh and collaborators in 2023 (24), entitled “*Wolbachia*-infected pharaoh ant colonies have higher egg production, metabolic rate, and worker survival”. From the dataset, the data used was from an experiment made in 2019 which compared egg-laying rates of queens in *Wolbachia*-infected and -uninfected colonies. Thirty-one colonies of *M. pharaonis* ants were observed and population data and egg laying rates were collected each three days, consisting of 46 days of experiment. Twenty colonies were infected with *Wolbachia*, and eleven composed the uninfected group. The experiment started when the queens were four days old.

The data downloaded from the Dryad was converted to an Excel document, having the following variables collected on the experiment:

- **date:** Date at which the data was collected
- **wolbachia:** Refers to *wolbachia* infection status.
- **day:** Number of days since the start of experimental protocol
- **colonyID:** Unique identifier for the colonies
- **q.age.days:** Age of queen(s) in days
- **males:** Number of males in the colony
- **workers:** Number of workers in the colony
- **queens:** Number of queens in the colony
- **eggs:** Number of eggs laid by the queens
- **yl:** Number of young larvae
- **ol:** Number of old larvae
- **worker.pp:** Number of workers pre-pupae
- **worker.pup:** Number of workers pupae

### Biodemography analysis

To verify the effect of the *Wolbachia* in the colonies using the formulas proposed by Birch, the variables ‘wolbachia’, ‘q.age.day’, ‘colonyID’, ‘queens’ and ‘eggs’ were selected for the analysis. For the calculation of the Intrinsic Rate of Increase (*r*), two rates were previously calculated, all with the use of the Excel software:

1. The age-specific survival rates (𝑙_𝑥_): in the case of this study, as all the colonies started with 20 queens, the 𝑙_𝑥_ was equal to the ‘nqueens’ divided by 20.
2. The age-specific fecundity rate (𝑚_𝑥_): equal to 0.5 multiplied by the ‘neggs’ and divided by the number of queens at the age x.

Given the reproductive and colony life type of ants (Hymenoptera: Formicidae), the ‘females’ were considered the queens only, besides the fact that ant workers are also female individuals, as the only reproductive ants are the queens.

Before the calculation of *r*, two important parameters of comparison were obtained, the Net reproduction rate (𝑅_0_), which is the ratio of total female births in two successive generations, and the Mean length of a generation (T). The 𝑅_0_can be calculated with the use of the following formula:

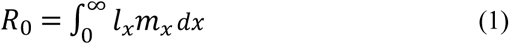

The 𝑙_𝑥_𝑚_𝑥_ factor was calculated before a simplification of the dataset. One value for each queen age was obtained for each variable by a simple mean of the values of the 31 different observed colonies. The 𝑅_0_was then calculated, along with the approximation of the Mean length of a generation (T), obtained with the following formula:

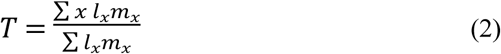

The calculation of the *r* was made with formula 3, for infected and uninfected colonies, using a procedure described in the article for the power of 𝑒. ‘𝑥’ here was used as the queen age in days.

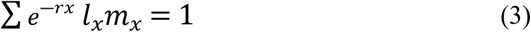

All the formulas were used for infected and uninfected colonies, for posterior comparison and analysis. Summing up, the analytical framework follows the life table approach pioneered by Lotka (1907) and Birch (1948), adapted to insect populations. Age-specific egg (“birth”) counts were transformed into fecundity rates (𝑚_𝑥_) and “survival” probabilities (𝑙_𝑥_), from which force of fecundity (𝜇_𝑥_) was derived. Empirical fecundity trajectories were then compared with classical parametric models explained in the following subsection.

### Classical demography

To detect fecundity deceleration and potential plateaus at later times, we estimated the logarithmic derivative of the force of mortality, the demographic measure known as the log-aging rate (LAR), defined as: 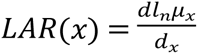.

Empirical LAR values were computed by age-specific differencing of log-transformed hazards, whereas model-based LARs were derived analytically from fitted hazard functions. Following approaches in biodemography (16,19), we defined the onset of fecundity deceleration as the first age at which LAR reached non-positive values for sustained consecutive ages. Bootstrap resampling was employed to assess uncertainty in estimates of the deceleration age. This work was done in R version 4.2.2 with a ‘Tidyverse’ library-based script. The dataset consisted of the life tables of both colonies (Tables 2 and 3). The values of *r* were calculated for posterior comparison with the Birch methodology described in 2.2.

The life-table aging rate (LAR) was obtained based on Horiuchi and Coale’s (1990) formula (4), where 𝑏(𝑥) represents the LAR and 𝑀(𝑥) is the central death rate at age 𝑥 (25)

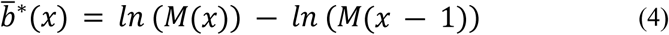

To avoid high fluctuations caused by variations in the death rates of older ages, Horiuchi and Coale proposed the empirical formula (5), which takes nine-year moving averages on the results of (4), after applying a five-year moving average to µ(𝑥).

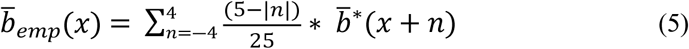

Nevertheless, due to the low number of observation days, this second-step smoothing procedure was unable to be employed.

To quantify fecundity patterns and detect late-life fecundity deceleration in ants, we combined empirical life table construction with parametric mortality models. Our analysis was grounded in classical biodemographic approaches(11,15,26), while adapting them to colony-level insect data.

In the present work, we fit four parametric mean functions µ(𝑥) representing per-queen fecundity at age x. These models provide alternative characterizations of age-specific reproductive schedules, and their parameters are estimated via maximum likelihood under a Poisson framework.

We thus fit a Gaussian (bell-shaped) approach to the queen’s fertility:

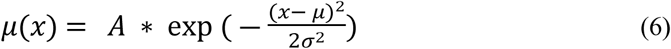

with parameters’ amplitude A ≥ 0, center μ, and spread σ > 0.

Next, a gamma Probability Density Function (PDF) was employed, with parameters 𝑘 𝑒 𝛾 where (𝑋 ∼ Γ(𝑘, 𝛾)):

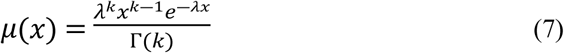

where, if 𝑘 = 1, the gamma distribution is reduced to the exponential distribution.

Third, we fit a Weibull PDF, with two positive parameters 𝑊𝑒𝑖𝑏𝑢𝑙𝑙 (𝜆, 𝛾):

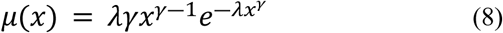

if 𝜆 = 1, we end up again with an exponential distribution, but with 𝜆 < 1the function increases monotonously, and if 𝜆 > 1, the opposite.

Lastly, we also employ a Gompertz–Makeham framework:

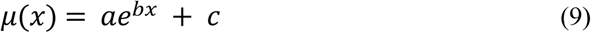

Parameters are also positive, where a ≥ 0, c ≥ 0. Here, the Gompertz–Makeham function – typically used for mortality – is repurposed to describe fecundity shapes, combining an exponential trend with a baseline floor.

All models were fitted following a Maximum Likelihood Estimation methodological approach where counts per age class 𝑦_𝑖_ are modeled as Poisson with mean 𝜆_𝑖_ = 𝐸_𝑖_µ(𝑥𝑖), where 𝐸_𝑖_ is exposure and µ(𝑥𝑖) is the model-predicted fecundity rate (log-likelihood):

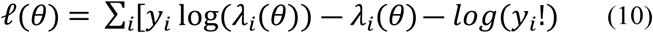

Additionally, model-based LAR was calculating using:

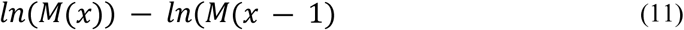

Evaluating plots include empirical LAR, smoothed trends (LOESS, GAM spline, cubic polynomial), and model-predicted LAR.

Comparisons were conducted across colonies to test for robustness of the observed patterns, while Residuals between empirical mortality rates and model predictions were computed to assess goodness-of-fit, and error distributions were analyzed to explore potential systematic deviations. This combined framework integrates classical demographic models with empirical hazard estimation, making it possible to rigorously test whether insect populations display fecundity plateaus analogous to those found in human and other animal demography.

### GEE modelling

To understand the influence of the presence of the bacteria and the time on the number of queens, eggs and workers, we used Generalized Estimating Equations (GEE), typical of marginal models (or population average models). This kind of model consider the non-independence of the observations, while not requiring assumptions about the distribution. While the linear models assume no correlation between the observations, the population average model provide a more robust inference and it does not specify any likelihood, being suitable for analyzing dependent observations, throughout time, between different treatments.

Although it is a useful analytical tool for evaluating relationships between different variables, Generalized Linear Models (GLM) can only be used when there is no correlation among the data. The GEE method uses quasi-likelihood methodology to create equations that link marginal means to linear predictors to provide a variance-covariance matrix (27).

As mentioned, GEEs are based on GLMs, so they also assume that the dependent variable belongs to the exponential family, but they include a correlation structure in the estimation of the regression parameters 𝜷 = [*β*_0_, *β*_1_, *β*_2_, …, *β*_*p*_].

The proposal by Liang & Zeger (28) consisted of solving the following equation to obtain the parameter estimates 𝜷

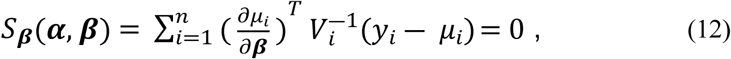

where 𝑉_𝑖_ is a specific matrix that incorporates a given correlation structure, and 𝜶 represents the parameters of that correlation structure,

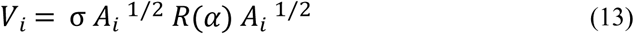

where 𝐴_𝑖_ is a diagonal matrix of the outcomes’ variance, 𝑅(𝛼) is the correlation matrix specified and σ is a dispersion parameter.

Liang & Zeger (28) demonstrate that if the structure for the mean is correctly specified, then, regardless of the chosen 𝑉 matrix, the estimator 𝜷 is unbiased and consistent. The distribution approaches to a multivariate normal distribution, that is

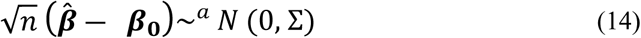

where Σ is the empirical estimation of variance. This property allows us to calculate p-values and confidence intervals using the Wald test.

The equations used in the correlation matrix are extensions of those used in the quasi-likelihood method. These likelihood methods are not Maximum Likelihood, since the distribution of Y is not specified, and therefore there is no likelihood function; the alternative computational method for clustered data is a generalization of quasi-likelihood (27)

The GEE method uses the quasi-likelihood methodology to create equations that link marginal means to linear predictors in order to provide a variance–covariance matrix (27). Therefore, GEEs do not require an assumed distribution for the response variable, provide more robust inference results, and do not specify any likelihood (29) In the present study, the observations made colony by colony form a dependent structure, since although the colonies themselves are independent, each one—considered an individual—was observed over time. Therefore, despite being widely used in biological and ecological studies, the use of GLM is not appropriate here, as they assume independence between observations and would not yield robust or reliable results for the *M. pharaonis* population and its relationship with infection by the endosymbiont *Wolbachia*. As the dependent variables analyzed consisted of counts, the distribution used was ‘Poisson’, which link function is the logarithmic, and the correlation structure considered was autoregressive of the first order (AR (1)). The analysis was done in the R software (version 4.2.2) (30) with the packages ‘geepack’, ‘dplyr’ and ‘ggplot2’.

## Results and discussion

### Descriptives of the data

The variables ‘wolbachia’, ‘day’, ‘colonyID’, ‘queens’, ‘eggs’ and ‘workers’ were selected for the analysis done in this work, as they are the variables related to the biodemography formulas used and describe the reproduction rates of the ant colonies, as well as the infection status by the *Wolbachia* bacteria. Since the data is not normally distributed and the variances between the infected and not infected group are not equal, the non-parametrical test of Mann-Whitney was made for each caste of the colony.

The mean number of queens infected with *Wolbachia* was 19.04, and the uninfected was 17.69 (Fig 3). The Mann-Whitney test showed that we can reject the null hypothesis that these means are equal (p<0.00).

**Figure 3:**
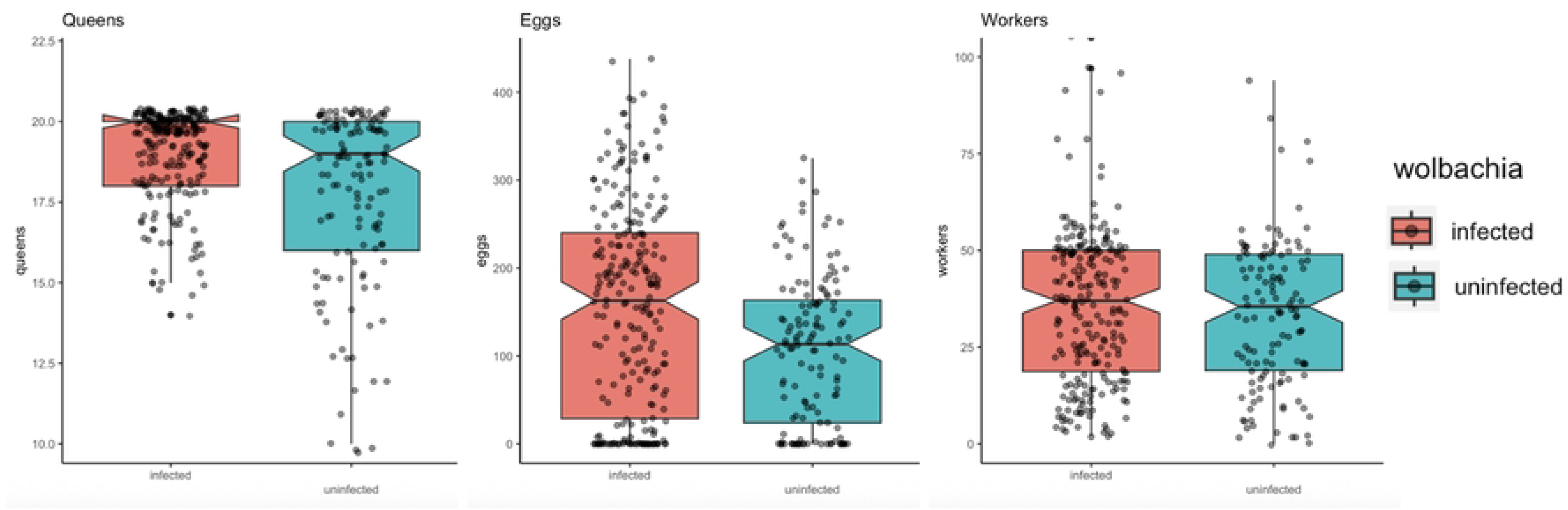
Boxplot of the number of queens, eggs and workers, respectively, in the infected (orange, left) and uninfected (green, right) colonies. Queens and eggs showed significant differences between treatments, while worker numbers showed no difference between treatments.

The mean number of eggs in the infected colonies with *Wolbachia* was 156, and in the uninfected was 107 (Fig.3). The Mann-Whitney test showed that we can reject the null hypothesis that these means are equal (p<0.00). The mean number of workers in the infected colonies with *Wolbachia* was 39.1, and in the uninfected was 39.7 (Figure 3). The Mann-Whitney test showed that we do not reject the null hypothesis that these means are equal (p=0.536).

### Biodemographic analysis

The mean number of queens and eggs per day of observation and the age-specific survival (𝑙_𝑥_) and fecundity (𝑚_𝑥_) rates were summarized in Table 1. The t test made with the number of queens from infected and uninfected colonies showed that the number of queens is significantly different between these two groups (p-value = 0.04629), but not the number of eggs laid by the queens (p-value = 0.2003).

**Table 1:**
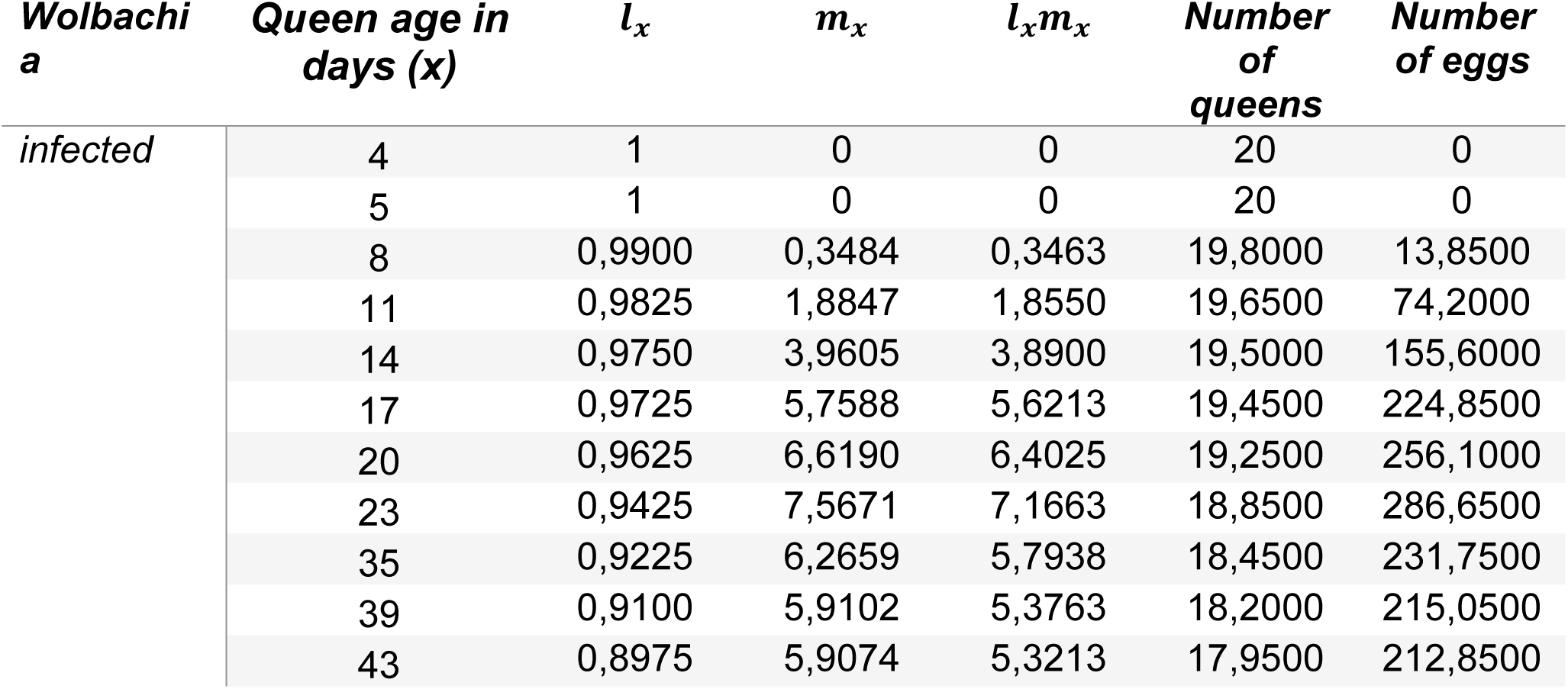
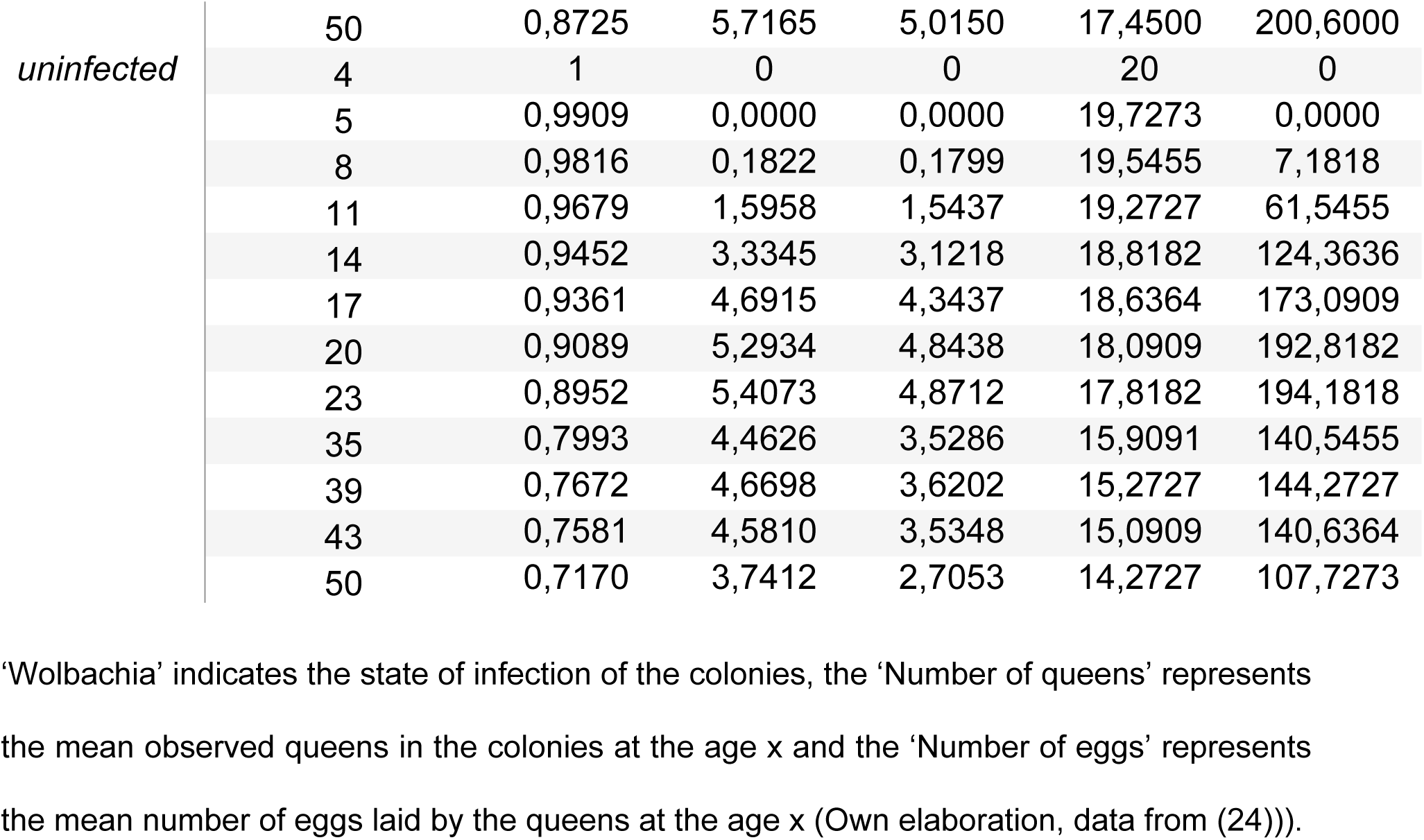
Simplified life table with the mean of each observation and the age-specific survival and fecundity rates for the infected ad uninfected colonies.

The calculation of the 𝑅_0_, by the sum of the values of 𝑙_𝑥_𝑚_𝑥_, showed different values for the two colony types. While the infected ones had 𝑅_0_ = 47, the uninfected ones had a 𝑅_0_= 32,29. This means that the presence of the *Wolbachia* bacteria makes the capacity the *M. pharaonis* colonies must grow in each generation go from 32,29 times to 47 times its original size. The values of T also favor the presence of *Wolbachia*, increasing the mean length of a generation by 1,31 days (from 27,71 to 29,02 days). The infinitesimal rate of increase, *r*, of the infected colonies (*r*= 0,18) was also higher when compared to the uninfected ones (*r*=0,16).

### Classical demography insights

Following the methodology specified in sections 3.2 and 3.3, it was possible to calculate diverse parameters trying to explain the evolution on the average number of eggs produced by the queen each day, i.e., queens’ fecundity. This analysis is possible to differentiate by infected and non-infected colonies. The next figures present an example of 8 random ant colonies: 4 infected and 4 uninfected. From Fig 4, on overall it is possible to identify a peak in the average number of eggs by queen in the middle of the observed time. Nevertheless, the fecundity of infected colonies is much higher than for uninfected.

**Figure 4:**
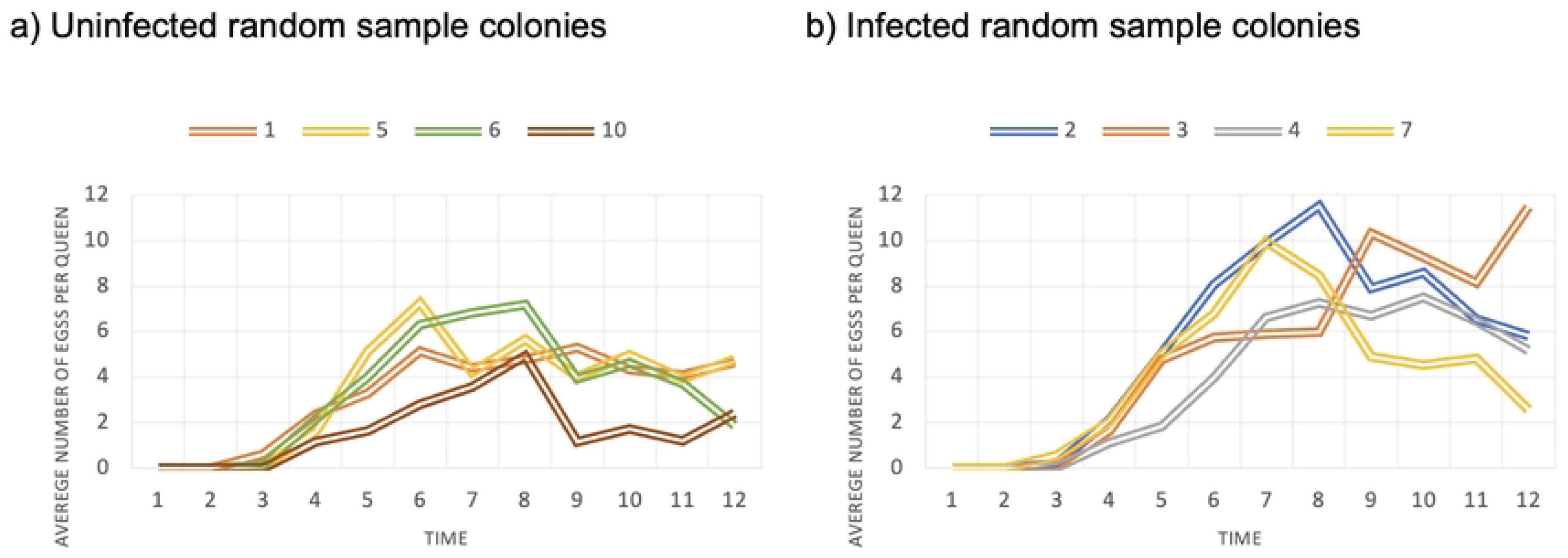
Average fecundity of queens over time. The infected colonies have a higher average number of eggs per queen, but its peak number occurs in older queens when compared to the uninfected colonies. Source: Own calculation.

Combined with the approaches exposed in equations (6) to (9), we can also try to find a proximity between laws of human demographic behavior and insect reproduction, in this case: ant colonies. Fig 5, presents, as example, empirical LAR and model fitting for an infected (labeled as colony “A”) colony a an uninfected (labeled as colony “U”) one.

**Figure 5:**
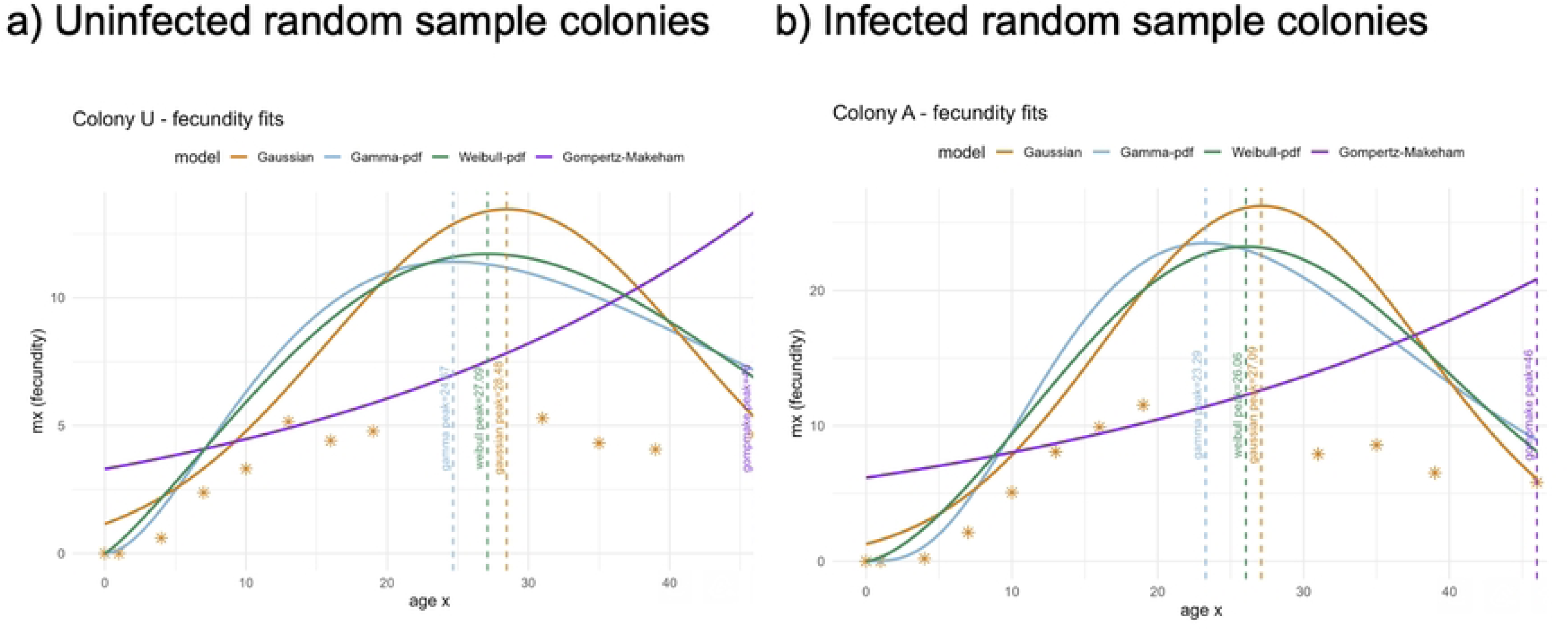
Life Table Aging Rate (empirical and theoretical). The models don’t follow closely the empirical LAR, but the curves show that the age which the fecundity starts to decelerate happens earlier in infected colonies. Source: Own calculation.

The presented results show that there is some difficulty to follow closely the empirical LAR (dots) but the two main models that visually present better results are the Weibull and the Gaussian. The left axis still shows the empirical differences in fecundity, with infected colonies presenting higher values, but the age at which fecundity starts to decelerate is later for uninfected ones.

In Fig 6, we can now confirm which of the models produces more accurate results. As we expected from Fig 5 observation, it is possible to conclude, for the presented examples, the gamma model is the one with lower residuals, i.e., the differences calculated between observed and estimated LAR values (𝑟_𝑖_ = 𝑦 ― 𝑦).

**Figure 6:**
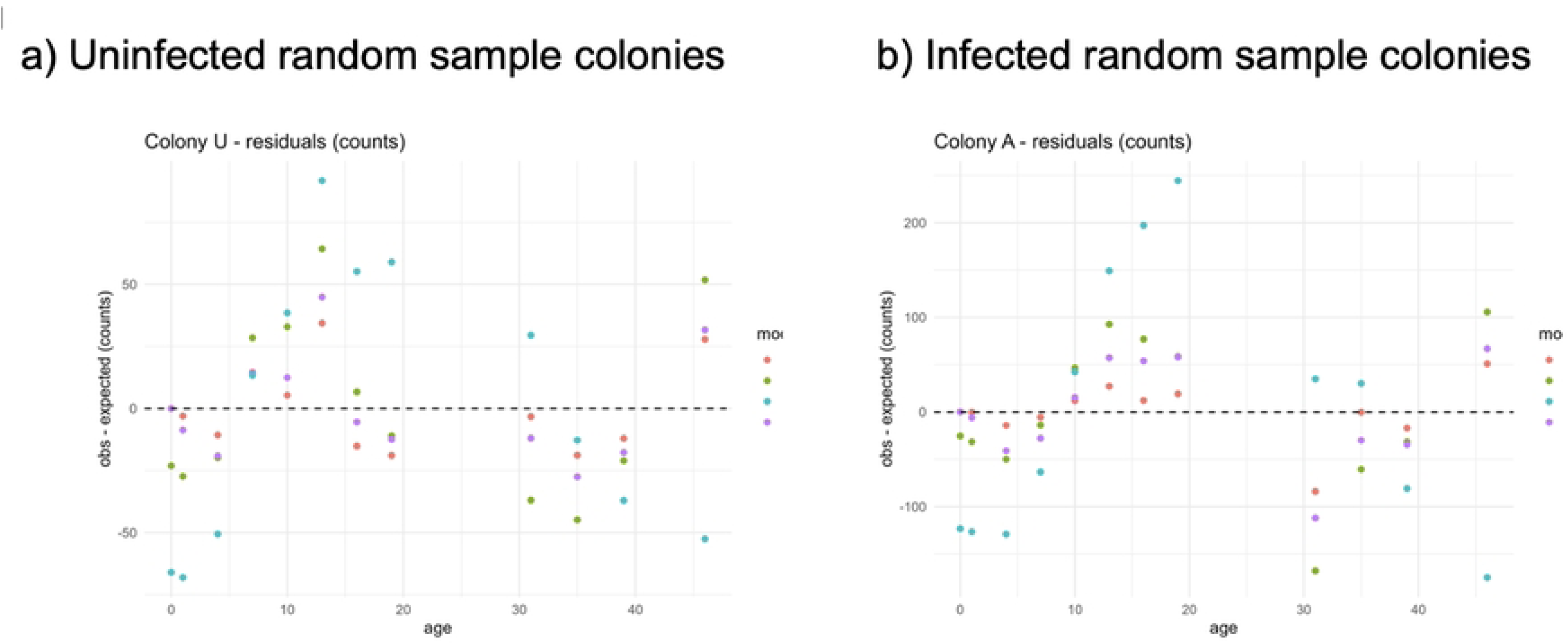
Fecundity Residuals (empirical - theoretical). The residual analysis show that the Gamma model has the lowest residuals. Source: Own calculation.

The gamma model estimates the age of fecundity deceleration for both colonies (infected and uninfected) a little lower than the Weibull and Gaussian Models. Nevertheless, our sensibility shows that for colony A (Fig 5 – infected) fecundity keeps increasing until age/day 20, and in colony U (uninfected) starts at day 13. So, besides if the gamma model estimates better the fecundity behavior, the Weibull and Gaussian models estimate more accurate the age at which deceleration starts.

On other hand, applying equation (4) it is possible to identify properly the time at which fecundity starts to decelerate. Thus, Fig 7 presents the Horiuchi Life Table Aging rate, but very conditioned once that the second step smoothing procedure from the original model is not possible to implement. In this case, it is only possible to make a very general conclusion: fecundity deceleration starts much earlier than expected by fitting the models to fecundity curves (Fig 5). Lastly, but still very interesting and important conclusion, is that the Birch intrinsic growth rate is higher for infected colonies when compared with uninfected.

**Figure 7:**
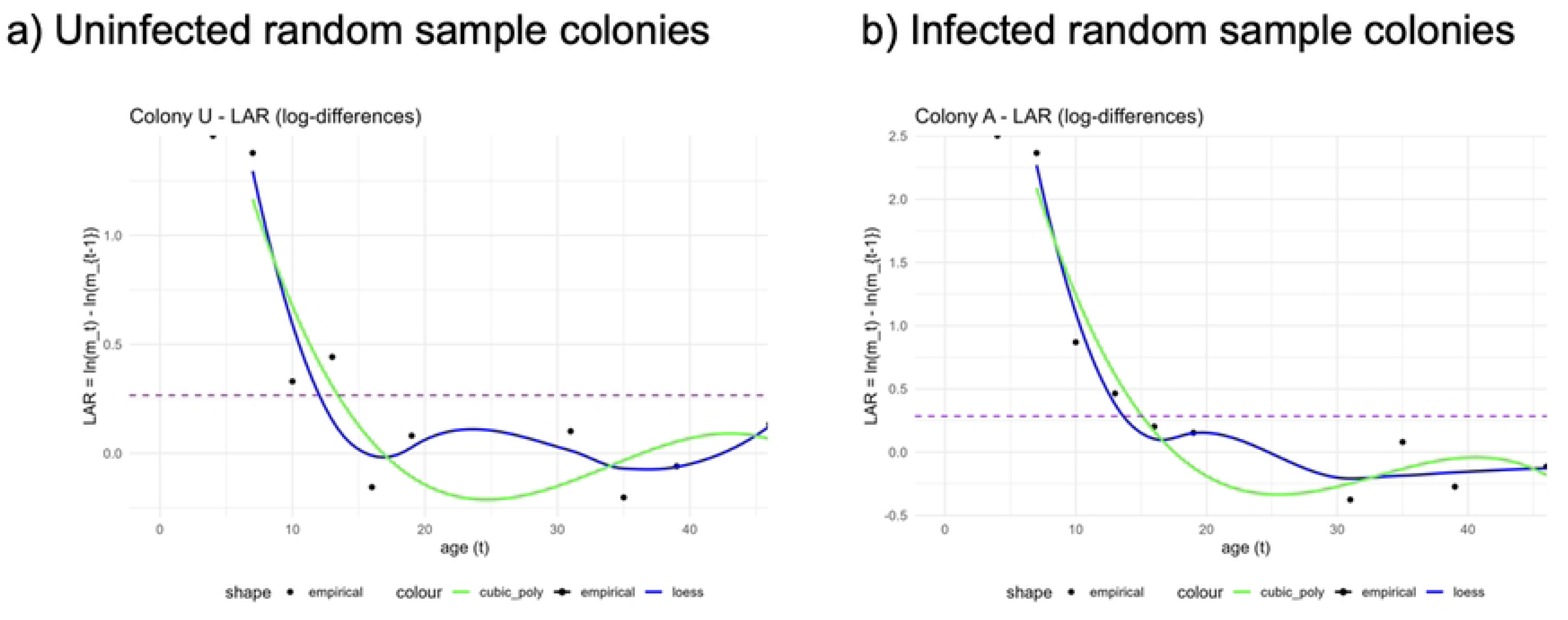
Empirical Life Table Aging Rate (observed and smoothed) and Birch intrinsic growth rate. By comparing the empirical LAR and the model fitting results, it is possible to see that the fecundity deceleration starts much earlier than expected by fitting the models to fecundity curves. The Birch Intrinsic Growth Rate, represented by the dotted line, is higher for the infected colonies. Source: Own calculation.

### GEE model analysis

One marginal model for each life stage used for the biodemography analysis was constructed, showing different results regarding the influence of the *Wolbachia* for each caste of the ants’ colony. The GEE models were constructed considering the family Poisson, with the logarithmic link function. The constraints of independence and AR(1) were tested and, since the coefficients were equal, we concluded that the correlation between the responses within the subjects is practically inexistent.

In order to build more realistic models, that take into consideration the effect of time in the *Wolbachia* infection in the colonies, a term of interaction between time and the *Wolbachia* infection was included in the models. The coefficients and p values for the queens’ model are in Table 2.

**Table 2:** Summary of the GEE model for the queen number in infected and uninfected colonies.

| Coefficients | Estimate | p-value |
| --- | --- | --- |
| (Intercept) | 2,99 | < 2e-16 |
| day | -0,02 | < 2e-16 |
| Day*wolbachiauninfected | -0,004 | 5.7e-07 |

This was the only model that showed a significant interaction between time and the bacteria infection, meaning that, in uninfected colonies, the decrease in queen number is more pronounced than in the infected colonies, indicating that, for the queens, the state of infection has an influence in the queen decrease throughout time. Considering the coefficients of Table 2 and the state of infection by *Wolbachia* as 0= ‘Infected’, and 1= ‘Uninfected’, this model indicates that the exponential expected number of queens in the infected colonies on day 0 is exp (2,99) =19,89, and the exponential expected number of queens decreases in a rate of exp (-0,02), or 2%, each day in infected colonies, and exp (-0,024), or 2,3%, in uninfected colonies.

For the eggs, a life stage related to the reproductive potential of the colony, the coefficients and p values is in Table 3.

**Table 3:** Summary of the GEE model for the eggs number in infected and uninfected colonies.

| Coefficients | Estimate | p-value |
| --- | --- | --- |
| (Intercept) | 4,48 | < 2e-16 |
| wolbachiauninfected | -0,37 | 1.4e-06 |
| day | -0,02 | < 2e-16 |

Considering the coefficients of Table 3 and the state of infection by *Wolbachia* as 0= ‘Infected’, and 1= ‘Uninfected’, this model indicates that the exponential expected number of eggs decreases in a rate of exp (-0,02), or 2%, each day in infected colonies, and exp (-0,39), or 32,29%, in uninfected colonies.

The coefficients were significant at a 5% level and show a positive influence of the presence of the bacteria on the number of eggs, but the infected and uninfected colonies have the same effect throughout time. The assumptions of the model residuals were evaluated.

For the workers, a caste not directly related to the colony’s reproduction, the following results were obtained (Table 4):

**Table 4:** Summary of the GEE model for the worker number in infected and uninfected colonies.

| Coefficients | Estimate | p-value |
| --- | --- | --- |
| (Intercept) | 4,15 | < 2e-16 |
| day | -0,03 | < 2e-16 |

The coefficient for the *Wolbachia* presence on the colony did not show a statistical significance for the worker numbers for this ant population, which indicates that the state of infection of the colony does not have an influence in the worker numbers, which decrease naturally through time. This model indicates that the exponential expected number of workers in the infected colonies on day 0 is exp (4,15) = 63,43, and the exponential expected number of workers decreases 2,3% each day.

## Discussion

It is already known that *Wolbachia* infections in *M. pharaonis* colonies have an effect in some reproductive dynamics (8). Besides that, the data used in the present work was analyzed in the original study (9) only in the scope of the number of queens and eggs laid and the expected age of the queens. The effect of these bacteria in ants is not widely known, and Singh and collaborators’ (8,9) work was one of the first experiments with the Pharaoh’s ants and the *Wolbachia* effect in the colony’s life cycle and growth (Ramalho et al., 2021).

Here, formulas based on demography studies of Birch (13) were used to check, within the scope of the biodemography, the effect of *Wolbachia* in the length of the generations, the capacity the colony has to grow in each generation and, mostly, if it had an effect on the infinitesimal rate of increase. The effect of *Wolbachia* in each caste throughout the days was also studied with the GEE modelling, a novel methodology in biology and ecology studies, as well as the application of classical demography concepts for this analysis.

The use of biodemography formulas, especially the Intrinsic Rate of Increase (*r*), is not frequent in ant colonies growth’ studies, so far. Until 2009, the knowledge on the *M. pharaonis* demography was not existent, although it is essential for the study of the population dynamics of this invasive species (1). The results obtained from the original study of the data, as the ones obtained in the present work, favor the presence of the bacteria for the longevity of the colony (9). However, it shows that the survival probability is similar in some stages of the colony life and that there is a similar proportion of alive queens over time.

The use of the rates calculated based on the work of Birch (13) presented results related to the reproductive capacity each colony type has, and, because of that, the use of biodemography formulas could be a way of having more specific analysis of the effect on the capacity the colony has as a whole, and not only the queens, the workers and the eggs alone. When thinking about control methods for ants, the calculation of the intrinsic rate is an interesting approach, as it takes in consideration the number of reproductive individuals and its individual and group capacity of producing eggs in the conditions the colony is in. This is a valuable tool on quantifying different variables on the colonies’ growth and on comparisons between different groups/treatments, in the scope of thinking more environmentally friendly control methods.

The results obtained in the present work show that the presence of *Wolbachia* increase the mean length of the generation, as well as the capacity it has to grow in each generation. With that, it becomes clearer that, along with the higher intrinsic rate of increase, the bacteria may give a reproductive acceleration to the Pharaoh’s ants colonies, giving it a capacity to grow more in a shorter period of time. As previous studies have shown that the size of the colony may influence the queen-worker ratio (2), an important next step could be studying the relation between colony size and the presence of *Wolbachia*, calculating the ratios used in the present work.

An important approach that should be considered is the use of biodemography concepts for workers analysis. As Kramer and collaborators (12) have shown, the lifespan of the colony influences the individual lifespan of workers and its sizes in *Lasius niger* - another ant species- colonies. Workers are responsible for the colony dispersion and a key factor on the invasion capacity of *M. pharaonis*. Adding the workers factor on the biodemography analysis can bring a new light on the understanding of *Wolbachia* and its use as a control mechanism for the Pharaoh’s ants.

The demography analysis of the LAR for the ant colonies indicates that the *Wolbachia* infection, besides giving the colonies a later peak on the eggs-per-queen numbers, a higher intrinsic rate of increase, generation length and the capacity the colony has to grow in each generation, the queen’s fecundity starts to decrease in earlier ages. This may indicate that the bacterial infection can act as a reproductive ’boost’, leading to a rapid increase in the colony’s reproductive capacity, followed by a decline in fecundity at a younger age compared to the absence of *Wolbachia*. This knowledge may aid future studies on the *Wolbachia–Monomorium pharaonis* mutualism, as well as on control methods for invasive ants that maintain relationships with these endosymbiotic bacteria, by supporting the development of strategies targeting the most appropriate life stages of the colonies.

Still within the scope of demographic analyses, the fitting of queen number over experimental time points (t) and their ages in days (x) using different models, i.e., Weibull, Gamma, and Gompertz, represents an uncommon practice in studies on ant colony growth and reproduction. To date, the application of such models in myrmecological research has been limited to studies on ant foraging activity (31). Based on the results obtained in the present study, this methodology is believed to offer substantial added value to research on colony growth in different ant species, particularly by enabling more accurate estimates of the ages at which queen fertility begins to decline, an important factor in studies aiming to develop control methods for invasive ants. It is worth emphasizing that data collection must be continuous and conducted at regular, closely spaced intervals in order to ensure more reliable model estimates.

The original study of the data, as well as other reproduction and growth studies, was analyzed with Generalized Linear Mixed Models (GLMM) for analyzing the differences in egg laying through time. Here, using GEE models, accurate and robust results were obtained regarding the influence of *Wolbachia* in the numbers of queens, workers and eggs during the days of the experiment. The presence of *Wolbachia* influenced the numbers of eggs and queens, corroborating the findings of previous studies (8,9)but not the worker numbers, which could indicate another evidence that *Wolbachia* is related to the reproductive capacity of the ants’ colonies. The GEE model for the queens also showed a significant interaction between the bacteria infection and the days of the experiment. This type of interaction, as other aspects of the *Wolbachia* infection in ants, has not been well studied, especially in a separated way for each caste and, given the simplicity of the GEE model construction and analysis, this could be a way to start analyzing this interaction and increase the knowledge on this symbiosis.

The understanding of *Wolbachia* interactions still need to increase (7) As *M. pharaonis* is an invasive species, many studies focus on control strategies and an understanding of the resource allocation (2), giving studies that have the objective of understanding the relations between the colonies and the bacteria’s presence an importance for the biological control of the ant species. The suppression of *Wolbachia* in *M. pharaonis* invaded places as a control method should decrease the reproduction capacity of the colonies and, by that, diminish the use of pesticides and the environmental impact.

## Conclusions

The results obtained in the present work confirmed our initial hypothesis that the presence of the endosymbiont bacteria *Wolbachia* increased the reproductive rates of *M. pharaonis* colonies. Besides that, it is also important to highlight the importance of using rates and concepts that go beyond the classical ecology and biology indexes and formulas. The use of classical demography adapted concepts showed a more quantifiable and integrated analysis of the reproductive differences between the colony types and, by that, a clearer vision of the effects *Wolbachia* has on *M. pharaonis*. Besides that, more studies are needed to fully comprehend this symbiotic relation and more interdisciplinary works need to be done on this and other biology themes.

The infection of the bacteria had a different influence in each caste. For the queens, the infected colonies showed to have a smaller decrease in their number throughout time, and the age in which their fecundity starts to decrease is earlier when compared to uninfected colonies. For the eggs, the peak if the number of eggs per queen happens in older ages in infected colonies, which also have more eggs when compared to uninfected colonies. For the workers, the infection state did not show a significant influence on their number. Considering the colony as a whole, the infection of *Wolbachia* increased the capacity the colony has to grow in each generation, its length and the colonies’ intrinsic rate of increase.

The results of the three areas of this study (biology, demography and statistics) proved to be complementary in providing a more holistic understanding of the effects of the endosymbiotic bacteria on both the distinct castes and the colony as an integrated unit. The methodologies applied in this work should be replicated to other ant species also mutualist to *Wolbachia* and to other experimental settings (as the comparison between food treatments), to study invasive species control methods, or even develop conservation actions and understand mutualistic relations between *Wolbachia* and the Formicidae.

## Acknoledgements

L.M.Machado acknowledges the R&D unit MED – Mediterranean Institute for Agriculture, Environment and Development (https://doi.org/10.54499/UIDB/05183/2020;https://doi.org/10.54499/UIDP/05183/2020) and the Associate Laboratory CHANGE – Global Change and Sustainability Institute (https://doi.org/10.54499/LA/P/0121/2020).

## Data Availability Statement

All data analyzed in this study are available at the Dryad Digital Repository (https://datadryad.org), on the dataset originally published by Singh et al. (2023) under the title ’*Wolbachia-infected pharaoh ant colonies have higher egg production, metabolic rate, and worker survival*’ and can be accessed via https://datadryad.org/dataset/doi:10.5061/dryad.0zpc8672b.

